# Metagenomic Noncoding RNA Profiling and Biomarker Discovery

**DOI:** 10.1101/2020.09.27.315507

**Authors:** Ben Liu, Sirisha Thippabhotla, Jun Zhang, Cuncong Zhong

## Abstract

Noncoding RNA plays important regulatory and functional roles in microorganisms, such as gene expression regulation, signaling, protein synthesis, and RNA processing. Given its essential role in microbial physiology, it is natural to question whether we can use noncoding RNAs as biomarkers to distinguish among environments under different biological conditions, such as those between healthy versus disease status. The current metagenomic sequencing technology primarily generates short reads, which contain incomplete structural information that may complicate noncoding RNA homology detection. On the other hand, de novo assembly of the metagenomics sequencing data remains fragmentary and has a risk of missing low-abundant noncoding RNAs. To tackle these challenges, we have developed DRAGoM (<u>D</u>etection of <u>R</u>NA using <u>A</u>ssembly <u>G</u>raph fr<u>o</u>m <u>M</u>etagenomics data), a novel noncoding RNA homology search algorithm. DRAGoM operates on a metagenome assembly graph, rather than on unassembled reads or assembled contigs. Our benchmark experiments show DRAGoM’s improved performance and robustness over the traditional approaches. We have further demonstrated DRAGoM’s real-world applications in disease characterization via analyzing a real case-control gut microbiome dataset for Type-2 diabetes (T2D). DRAGoM revealed potential ncRNA biomarkers that can clearly separate the T2D gut microbiome from those of healthy controls. DRAGoM is freely available from https://github.com/benliu5085/DRAGoM.

## INTRODUCTION

Noncoding RNA (ncRNA) can perform versatile functional roles than just acting as a genetic information carrier, and its importance in cellular physiology is increasingly recognized. For example, riboswitch is a class of *cis*-regulator locates in the 5’UTR of its target gene and can alter the gene’s expression efficiency by alternating its fold structure upon the binding with small metabolites or ion ligands [1–3]. A different *trans*-regulatory mechanism was found to be exerted by bacterial small RNAs (sRNA), which in most cases attenuate their target mRNA expressions by sequence complementarity-based binding (in a similar way as eukaryotic microRNAs) [4–7]. Noncoding RNAs can also catalyze biochemical reactions (ribozymes) [8], as exemplified by the well-known ribosomal RNAs (which catalyze protein synthesis) and group I and II introns (which catalyze the excision of themselves from the transcript) [9, 10]. With the prevalence of metagenomics [11–16], more microbial genomics sequences, including the previously uncharacterized ones, have been deposited into public databases. The amazing richness of microbial genomic data renders an unprecedented opportunity for ncRNA study. Indeed, the diversity and richness of microbial ncRNA function revealed from analyzing metagenomic data are beyond our existing knowledge [17–20], including the discovery of many long noncoding RNA classes such as OLE, GOLLD, and HEARO, with exceptionally complicated structures [21]. These discoveries underpin the importance of ncRNA functions in bacterial physiology, ecology, and interaction with the environment.

Despite the importance of functional ncRNAs, their use as biomarkers to characterize environments under different biological conditions (e.g., disease vs. healthy) appeared to be less explored than the protein-coding genes. Currently, only few ncRNAs (such as 16s rRNA) have been used as proxies to infer the taxonomic compositions of microbial communities [22–28]. On the other hand, systematic analyses of a broad range of protein-coding gene families have been routine in metagenomics. The analyses typically include the quantification of bacterial genes and gene families, the reconstruction of metabolic pathways, and the development of mathematical models to characterize microbiomes from different environments [29, 30]. For example, an increased membrane sugar transportation activity was associated with Type-2 diabetes (T2D) [31], a more active lactate metabolism may enhance athlete performance [32], and the dysfunction of nitrotoluene degradation may cause pediatric Crohn’s disease [33]. In addition to functional insights, protein marker gene analyses also offer a higher resolution in taxonomic profiling than using 16s rRNA alone [34]. Given the success of functional gene analysis in metagenomics, it is tempting to extend the analysis to functional ncRNAs and explore their potential as biomarkers.

Similar to protein, the key to the success of ncRNA biomarker discovery is the homology search. That is, whether we can reliably detect ncRNA reads from metagenomic sequencing datasets and assign them into different ncRNA families. Because the function of ncRNA is determined by both its structure and primary sequence (in few cases by primary sequence alone, such as microRNA [35–37]), the homology search of ncRNA often relies on the conservation of both types of information [38, 39]. They can be further compiled into a covariance model (CM) using stochastic context-free grammar (SCFG) to characterize a given ncRNA family [40], in a similar way as using profile hidden Markov model (profile-HMM) for protein family representation [41]. In the context of metagenomic sequencing data, the short reads (~100-150bp) may only contain partial secondary structure information, leading to inferior ncRNA homology search performance. The issue has been partially addressed by the development of the trCYK algorithm for homology detection with incomplete secondary structure [42], but its performance remained lower than directly operating on complete genomes. On the other hand, while a natural way to resolve this issue is to reconstruct complete secondary structure information via de novo genome assembly, assembly itself remains fragmentary and challenging [43–46]. Many ncRNA reads, especially the low-abundant ones, may not be assembled into contigs and are therefore missed in the subsequent homology stage.

To tackle the challenge of ncRNA homology search from metagenomic sequencing data, we have developed DRAGoM (<u>D</u>etection of nc<u>R</u>NA on <u>A</u>ssembly <u>G</u>raph <u>o</u>f <u>M</u>etagenomic data). DRAGoM is the first ncRNA homology search method that operates on sequence assembly graph, which sets it apart from the traditional methods that operate on unassembled reads (hereafter referred to as the “read-based” strategy) or assembled contigs (hereafter referred to as the “assembly-based” strategy). Homology search on assembly graph has been proven successful for protein [47–49] and protein family [50], and we expect to extend the success to ncRNA. Specifically, note that a path in an assembly graph corresponds to a set of overlapping reads, which may contain more complete secondary structure information that facilitates homology detection. Hence, we can expect DRAGoM to outperform the read-based strategy. Meanwhile, using the complete set of paths in the assembly graph without topological simplification (e.g., bubble removal and tip trimming [51–53]) and traversal (e.g., as Eulerian paths [54]) is more likely to preserve the original metagenome information (such as polymorphism and stain-level sequence variation). As a result, DRAGoM could rescue some of the ncRNA reads that are difficult to be assembled and outperform the assembly-based strategy. The only concern would be computational efficiency. However, we show that with proper path filtering (details in the Materials and Methods section), DRAGoM can be used to profile more than one thousand ncRNA families on medium-complexity datasets (e.g., human gut microbiome).

We have benchmarked DRAGoM with the read-based strategy and the assembly-based strategy. We chose CMSearch [55], which has included the trCYK algorithm [42], as the representative of the read-based strategy. We selected the string graph assembler SGA [53] and the de Bruijn graph assembler SPAdes [51, 52] for the assembly-based strategy (their contigs were subsequently searched using CMSearch). Our benchmark experiment considered both simulated and real datasets involving the search of an extensive collection of Rfam [56] ncRNA families. DRAGoM showed a higher and most robust performance compared to the other competitors. DRAGoM also improved homology search of 16s rRNA. Finally, we applied DRAGoM on real metagenomic datasets from T2D patient and control samples, and successfully discovered many ncRNA biomarkers for characterizing the T2D gut microbiome. The results imply the potential clinical applications of ncRNAs as biomarkers for disease diagnosis/prognosis. DRAGoM is freely available from https://github.com/benliu5085/DRAGoM.

## MATERIALS AND METHODS

### The DRAGoM Algorithm

The DRAGoM algorithm contains two main stages: (1) the construction of a hybrid assembly graph and (2) the identification of homologous ncRNA elements from the resulting hybrid assembly graph. By hybrid assembly graph, we mean the assembly graph that results from merging a string graph [57] and a de Bruijn graph [58], the two main computational models used in sequence assembly. A string graph is constructed based on the suffix-prefix overlaps between reads, whereas a de Bruijn graph is constructed based on the shared fc-mers between reads. Each model has its advantages and limitations, with the string graph being comparably more accurate but fragmentary. Both models have been integrated to improve sequence assembly [59]. To illustrate the idea, we present a toy example in Figure 1(A). The top-left panel shows an artificial genome sequence and the corresponding short reads. The bottom-left panel shows the string graph constructed from the reads with a minimum overlap length of 4bp. Because of the lower coverage at the middle of the artificial genome, only four reads can be overlapped. A missing link (the blue dash line) exists between the two subgraphs, leading to a subsequent fragmentary assembly. For the de Bruijn graph construction (the top-right panel), all reads can be connected using 3-mers as the vertices. While the de Bruijn graph completely recovers all reads, its graph topology is complex and can be traversed in two ways (with or without going into the loop). However, note that one of these de Bruijn graph traversals (i.e., with the loop) can be aligned to two terminal edges in the string graph (bottom-right panel, underlined sequences). Once the corresponding string-graph edges were reconnected using the de Bruijn graph path, the resulted hybrid graph perfectly represented the original genome.

**Figure 1:**
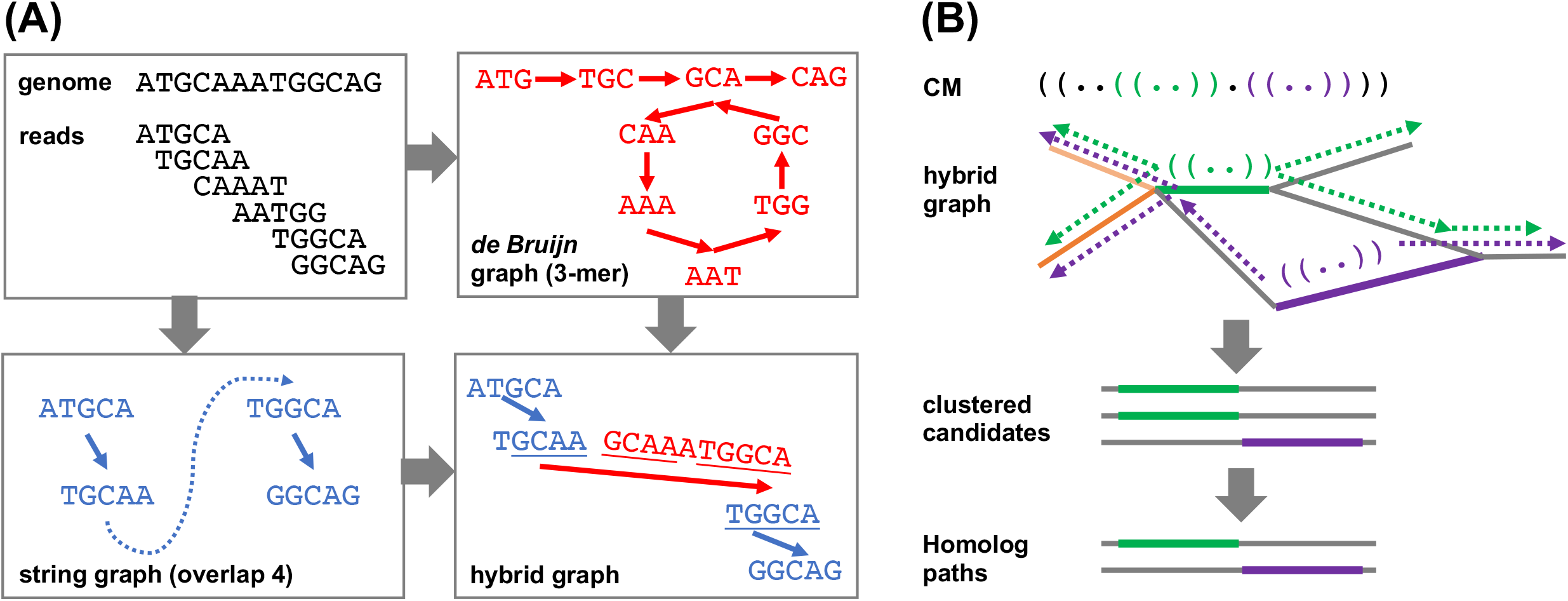
A schematic illustration of the DRAGoM algorithm. (A) The construction of the hybrid assembly graph. The hybrid graph, resulted from the merge of a de Bruijn graph and a string graph, can perfectly represent the original genome used in this toy example. (B) The search of ncRNA homologs against a hybrid graph. The green and purple parenthesis in the querying ncRNA CM (covariant model) represent local secondary structural components. The thick green and purple lines in the hybrid graph indicate anchors for path extension. Arrows indicate path extensions of the corresponding anchors.

The hybrid graph construction stage of DRAGoM implemented the above intuition. Specifically, SGA (version 0.10.15) [53] was used to generate the string graph, and SPAdes (version 3.13.0) [52] was used to generate the de Bruijn graph. When running SPAdes, the “--meta” tag was set to indicate metagenomic input (also known as “metaSPAdes”). Both programs were run in the paired-end mode. Detailed command lines for running both assemblers are available in the Supplementary Methods. The intermediate output of SGA (i.e., the .asqg file) was further simplified (using in-house scripts) to condense unbranched paths into single edges. Terminal edges (i.e., edges with in-degree or out-degree of 0) of the resulted string graph were then aligned with the trusted SPAdes contigs (no coverage hole, see more in Supplementary Methods) using BWA [60]. Only alignments with score >45 (per the BWA manual, +1 for a match, −4 for a mismatch, and −6 for a gap), alignment length >100, and no clipping at the open end (i.e., the end with a degree of 0 in the string graph) were considered. Then, for each SPAdes contig, if it had recruited more than one alignment, the corresponding terminals in the string graph defined by any pair of alignments were connected using the corresponding interval sequence of the SPAdes contig. If a SPAdes contig had recruited only one alignment, the corresponding string graph terminal was extended using the contig’s corresponding prefix or suffix. SPAdes contigs with no recruited alignment were also retained as isolated vertices in the hybrid graph. In a CAMI [44] dataset that contained ~15M vertices in the string graph, ~0.7M such connections were made.

The second main stage of the DRAGoM algorithm is to identify homologous paths of the querying CM from the resulted hybrid assembly graph. Intuitively, one can exhaustively enumerate all hybrid-graph paths and align them with the querying CM. However, this naïve approach would be practically infeasible because the number of paths grows exponentially with the number of reads in the dataset. We designed a filter-based heuristic for the speedup (Figure 1(B)). First, the querying CM was aligned to each hybrid-graph edge (which corresponds to a condensed path without branching). The edges bearing significant similarity to the querying CM were recorded as anchors. This stage allowed the detection of conserved short structural components (e.g., the green and purple stem-loops in the CM and the bolded paths in the hybrid graph of Figure 1(B)). The anchors were then extended towards both directions, aiming to reconstruct complete sequences of the candidate ncRNA homologs (the broken arrows in Figure 1(B)). The extension lengths for each anchor were determined by the unaligned prefix and suffix lengths of the CM (with a further extension of 10% length to account for potential gaps). Because some edges of the hybrid graph might represent similar sequences (e.g., the heavy and light orange edges in Figure 1 (B)), all paths resulted from extending the anchors were subject to sequence redundancy removal using CD-Hit [61]. Finally, the set of non-redundant paths were re-aligned to the querying CM, and the paths passing the gathering score threshold were selected as homologs of the corresponding ncRNA family. Individual reads were further mapped to the homologous paths for their functional annotations and the quantification of the corresponding ncRNA families. More details regarding this stage can be found in Supplementary Methods.

The above algorithm was implemented as the DRAGoM software package. DRAGoM accepts a set of querying CM and a given metagenomics sequencing dataset; it outputs the ncRNA read annotation and the quantification of each querying ncRNA family. DRAGoM was implemented using GNU C++ and Python and has been tested under several major Linux distributions (RedHat, Fedora, and Ubuntu). It is freely available from https://github.com/benliu5085/DRAGoM under the Creative Commons BY-NC-ND 4.0 License Agreement (https://creativecommons.org/licenses/by-nc-nd/4.0/).

### Benchmark Datasets

We have used six datasets to benchmark the performance of DRAGoM (Table 1). Detailed information regarding the reference genomes included in the community, their respective relative abundances, and the in-silico simulation command lines is available from Supplementary Table 2. All datasets, except one that can be directly downloaded from NCBI SRA, are available from https://cbb.ittc.ku.edu/DRAGoM.html. These six benchmark datasets include:

- *DS1* (*the REAGO dataset*): This simulated dataset represented a low-diversity metagenomic dataset that contains microbes from different clades with staggered abundances. The dataset was used in the benchmark experiment of REAGO [24]. It was simulated in silico with an average read length of 100nt and an expected error rate of 1%, containing 4,653,918 paired-end reads.
- *DS2* (*the Streptococci dataset*): This simulated dataset represented a community with highly-related microbial genomes from the same genus (e.g., *streptococcus*). The dataset was simulated in silico using eight streptococcus genomes, with an average read length of 100nt and an expected error rate of 1%. This dataset contained 600,000 paired-end reads.
- *DS3* (*the marine dataset*): This dataset represented a subset of microbial metagenome that was frequently observed from the marine environment. It was simulated from 28 marine genomes with an average read length of 100nt and an expected error rate of 1%, resulted in 3,700,000 paired-end reads.
- *DS4* (*the subsampled gut dataset*): This dataset represented a real human gut microbiome community (SRR341583). To facilitate the generation of ground-truth homology for benchmarking, we subsampled the dataset via read mapping against a set of microbial genomes that were frequently found in human gut. Only reads mapped to the selected reference genomes were retained, leaving 11,228,362 paired-end reads for this dataset.
- *DS5* (*the subsampled CAMI dataset*): This dataset was downloaded from CAMI [44], a comprehensive simulated dataset. To focus on the more challenging cases of metagenomics analysis, only reads representing low-coverage genomes (<10X) were selected (via read mapping). This dataset contained 31,311,294 paired-end reads.
- *DS6* (*the T2D dataset*): This dataset contained twelve samples (six cases and six controls) generated for the study of Type-2 Diabetes (T2D) gut microbiome. Specifically, the samples SRR341616, SRR341617, SRR341618, SRR341621, SRR341623, and SRR341624 were used as the healthy controls; and the samples SRR341583, SRR341584, SRR341585, SRR341586, SRR341587, and SRR341588 were used as T2D cases. To eliminate sex bias, all samples selected came from female donators. All datasets contain paired-end reads, with sizes ranging from 42M-57M (after quality trimming). They were selected to demonstrate the practical applications of DRAGoM.

**Table 1.** Summary of experimental datasets.

| Dataset | Description | No. of Genomes | Abundance | No. Reads | Read Length | Error rate |
| --- | --- | --- | --- | --- | --- | --- |
| DS1 | REAGO | 14 | Staggered | 4.6M | 100 | 1% |
| DS2 | Streptococcus | 8 | 5x | 0.6M | 100 | 1% |
| DS3 | Marine | 28 | 5x | 3.7M | 100 | 1% |
| DS4 | Human gut | 3,499 | Staggered | 11.2M | 74 | - |
| DS5 | CAMI | 4,679 | Staggered | 31.3M | 100 | - |
| DS6 | T2D | - | - | - | 74 | - |

### Benchmark Experiment Setup

Given a querying ncRNA family, we define its ground-truth homologs as the reads that can be generated or mapped (>60% of their total lengths) to the genomic intervals that were annotated as the ncRNA family by CMSearch [55] (under its default stringency cutoff). The command lines used for ground-truth generation are available from Supplementary Methods.

Given the ground-truth definition, we defined true positives (TP) as the identified homologous reads. We further defined false positives (FP) as the identified non-homologous reads, and false negatives (FN) as missed homologous reads. We further defined recall and precision as:

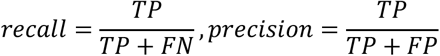

and subsequently F-score as:

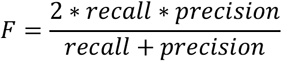

All methods were tested under various stringency cutoffs to generate the receiver operating characteristic (ROC) curve. The ROC curves were extrapolated to the points (recall: 0, precision: 1) and (recall: 1, precision: 0) to calculate the area under the curve (AUC).

We benchmarked our graph-based ncRNA homolog search strategy DRAGoM (homolog search against assembly graph) with the read-based strategy (homolog search against unassembled reads) and the assembly-based strategy (homolog search against assembled contigs). For the read-based strategy, we chose CMSearch as the representative; we refer it as “CMSearch” hereafter. For assembly-based strategy, we chose SGA (as the representative of string graph assemblers) and SPAdes (as the representative of de Bruijn graph assemblers); hereafter we refer them as “SGA+CMSearch” and “SPAdes+CMSearch”, respectively. Command lines for executing the programs are available in Supplementary Materials. Each method was benchmarked using different sets of querying ncRNA families (details available from Supplementary Table 2). The reported performance corresponded to the unweighted arithmetic mean among the set of querying ncRNA families. Note that the performance for 16s rRNA was reported individually, given its importance in taxonomic profiling.

## RESULTS

### Performance on Simulated Datasets

The performances of all tested methods on DS1 (the REAGO dataset, 42 ncRNA families searched) are shown in Figure 2. For non-16s rRNA queries (Figure 2(A)), DRAGoM was able to achieve the highest recall, representing a gain of 7.3% recall rate as compared to the second-best performer SPAdes+CMSearch (Table 2). CMSearch alone performed significantly worse than DRAGoM and SPAdes+CMSearch, potentially due to the lack of complete secondary structure information in unassembled reads. SGA+CMSearch seemed to be adversely impacted by the low coverage of this dataset and showed the lowest recall. However, it did show the highest precision rate. The observation was in line with the difference between string graph and de Bruijn graph assemblers observed elsewhere. In terms of the peak F-score, DRAGoM achieved 93.6%, followed by 92.2% of SPAdes+CMSearch. In terms of AUC, DRAGoM was also the best performer with 96.8%, compared to 93.9% of the second-best method SPAdes+CMSearch. For 16s rRNA, all methods performed well (Figure 2(B)). DRAGoM remained the best method with a marginal improvement (99.5% F-score and 99.6% AUC, followed by 97.6% F-score and 98.8% AUC of the second-best method CMSearch, see Table 4). Surprisingly, SPAdes+CMSearch showed the lowest sensitivity, potentially due to the polymorphism information lost during the graph simplification and traversal stages of SPAdes. Overall, DRAGoM showed a higher performance than any tested method and was robust for both non-16s and 16s rRNA searches.

**Figure 2:**
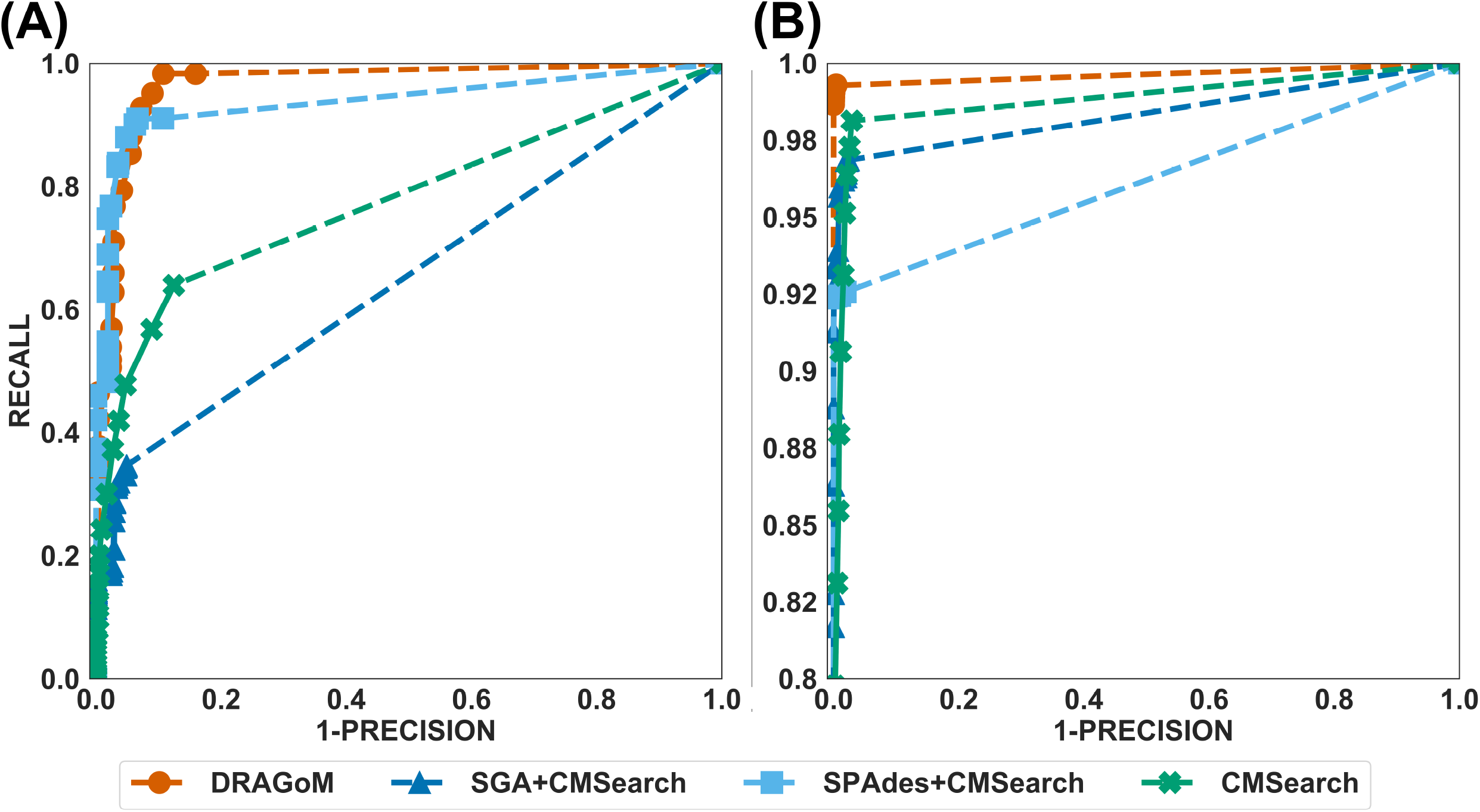
The ROC curves for searching (A) 42 non-16s rRNA ncRNA families and (B) 16s rRNA using the corresponding programs against DS1 (the REAGO dataset).

**Table 2.**
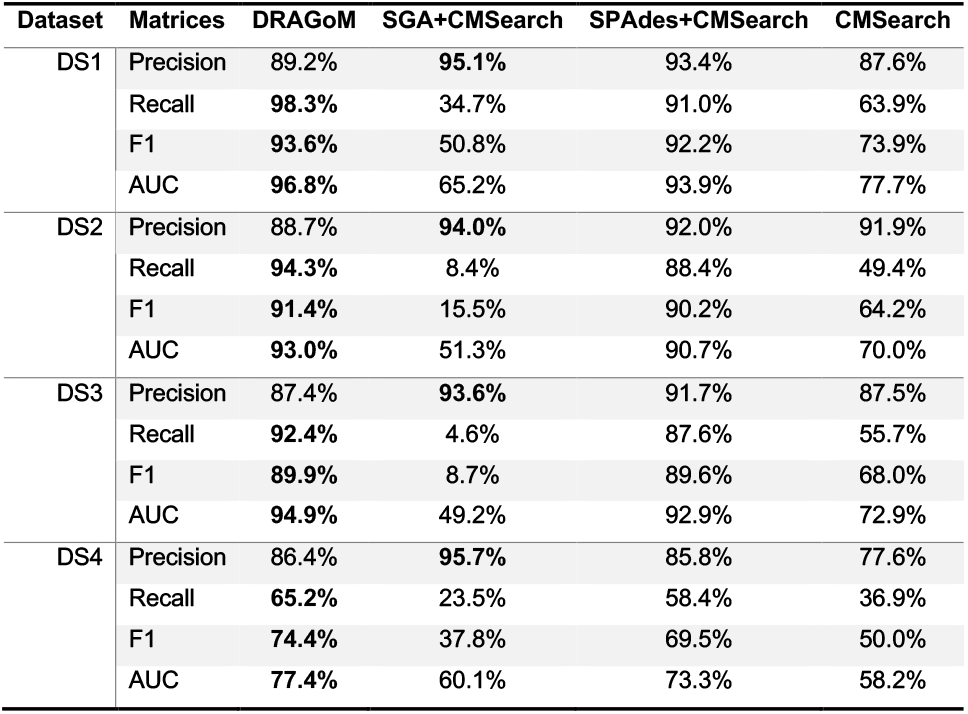
Performance summary of the tested methods on DS1-DS4 (for non-16s rRNA queries). The highest performance of each category is bolded.

| Dataset | Matrices | DRAGoM | SGA+CMSearch | SPAdes+CMSearch | CMSearch |
| --- | --- | --- | --- | --- | --- |
| DS1 | Precision | 89.2% | **95.1%** | 93.4% | 87.6% |
|  | Recall | **98.3%** | 34.7% | 91.0% | 63.9% |
|  | F1 | **93.6%** | 50.8% | 92.2% | 73.9% |
|  | AUC | **96.8%** | 65.2% | 93.9% | 77.7% |
| DS2 | Precision | 88.7% | **94.0%** | 92.0% | 91.9% |
|  | Recall | **94.3%** | 8.4% | 88.4% | 49.4% |
|  | F1 | **91.4%** | 15.5% | 90.2% | 64.2% |
|  | AUC | **93.0%** | 51.3% | 90.7% | 70.0% |
| DS3 | Precision | 87.4% | **93.6%** | 91.7% | 87.5% |
|  | Recall | **92.4%** | 4.6% | 87.6% | 55.7% |
|  | F1 | **89.9%** | 8.7% | 89.6% | 68.0% |
|  | AUC | **94.9%** | 49.2% | 92.9% | 72.9% |
| DS4 | Precision | 86.4% | **95.7%** | 85.8% | 77.6% |
|  | Recall | **65.2%** | 23.5% | 58.4% | 36.9% |
|  | F1 | **74.4%** | 37.8% | 69.5% | 50.0% |
|  | AUC | **77.4%** | 60.1% | 73.3% | 58.2% |

For DS2 (the Streptococcus dataset, 27 ncRNA families searched), all methods behaved similarly as in DS1 for non-16s searches (Figure 3(A)). DRAGoM again performed the best on this dataset (91.4% F-score and 93.0% AUC), followed by SPAdes+CMSearch (90.2% F-score and 90.7% AUC, see Table 2). The lower performances of CMSearch and SGA+CMSearch were also observed as in DS1, and may due to similar reasons as discussed previously. For 16s rRNA (Figure 3(B)), SGA+CMSearch performed the best (99.2% F-score and 99.8% AUC), with DRAGoM as the second-best method in F-score (98.1%) and CMSearch in AUC (99.4%, see Table 4). SGA+CMSearch seemed to benefit from its preservation of polymorphism information in 16s rRNA by adopting a more conservative graph simplification strategy. On the other hand, DRAGoM remained the most sensitive method (with the highest recall rate of 99.9%), but its overall performance appeared to be compromised by the lower precision rate due to exhaustive path traversal (96.2%, see Table 4).

**Figure 3:**
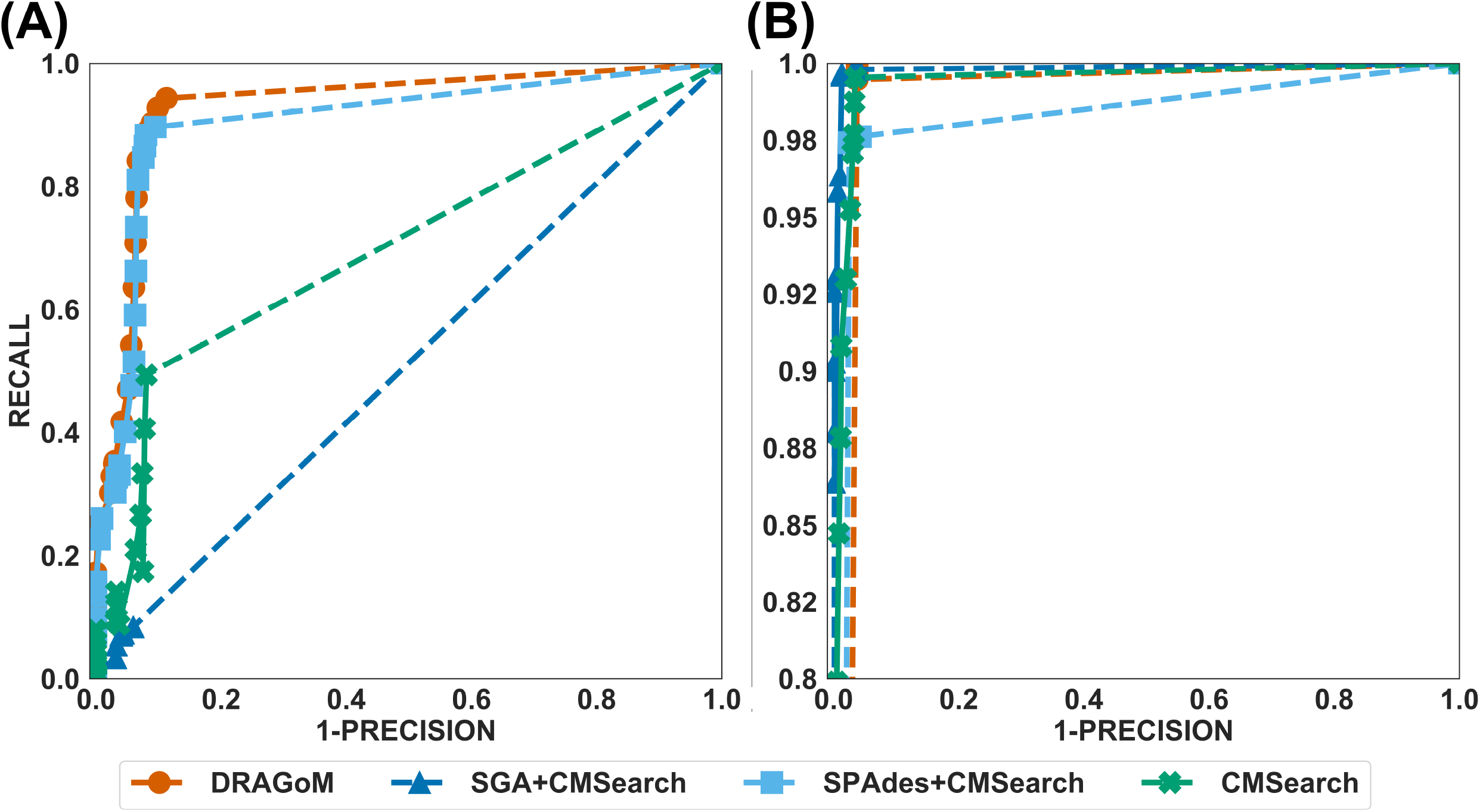
The ROC curves for searching (A) 27 non-16s rRNA ncRNA families and (B) 16s rRNA using the corresponding programs against DS2 (the simulated Streptococcus dataset).

For DS3 (simulated marine, 93 ncRNA families searched; see Figure 4) and DS4 (subsampled human gut, 60 ncRNA families searched; see Figure 5), these methods also performed similarly as in DS1 and DS2. DRAGoM outperformed the other methods in non-16s rRNA queries, reaching 89.9% F-score and 94.9% AUC for DS3 (Figure 4(A)), and 74.4% F-score and 77.4% AUC for DS4 (Figure 5(A)). The lower performance on DS4 because it is a real dataset that contains more experimental noise than the simulated ones. SPAdes+CMSearch also remained as the second-best method on both DS3 and DS4. For 16s rRNA, DRAGoM performed the best on DS3 (99.1% F-score and 99.3% AUC; see Figure 4(B) and Table 4). On DS4, SGA+CMSearch performed the best (96.1% F-score and 96.4% AUC; see Figure 5(B) and Table 4), followed by DRAGoM (94.2% F-score and 94.4% AUC). These observations were also consistent with those made in DS1 and DS2.

**Figure 4:**
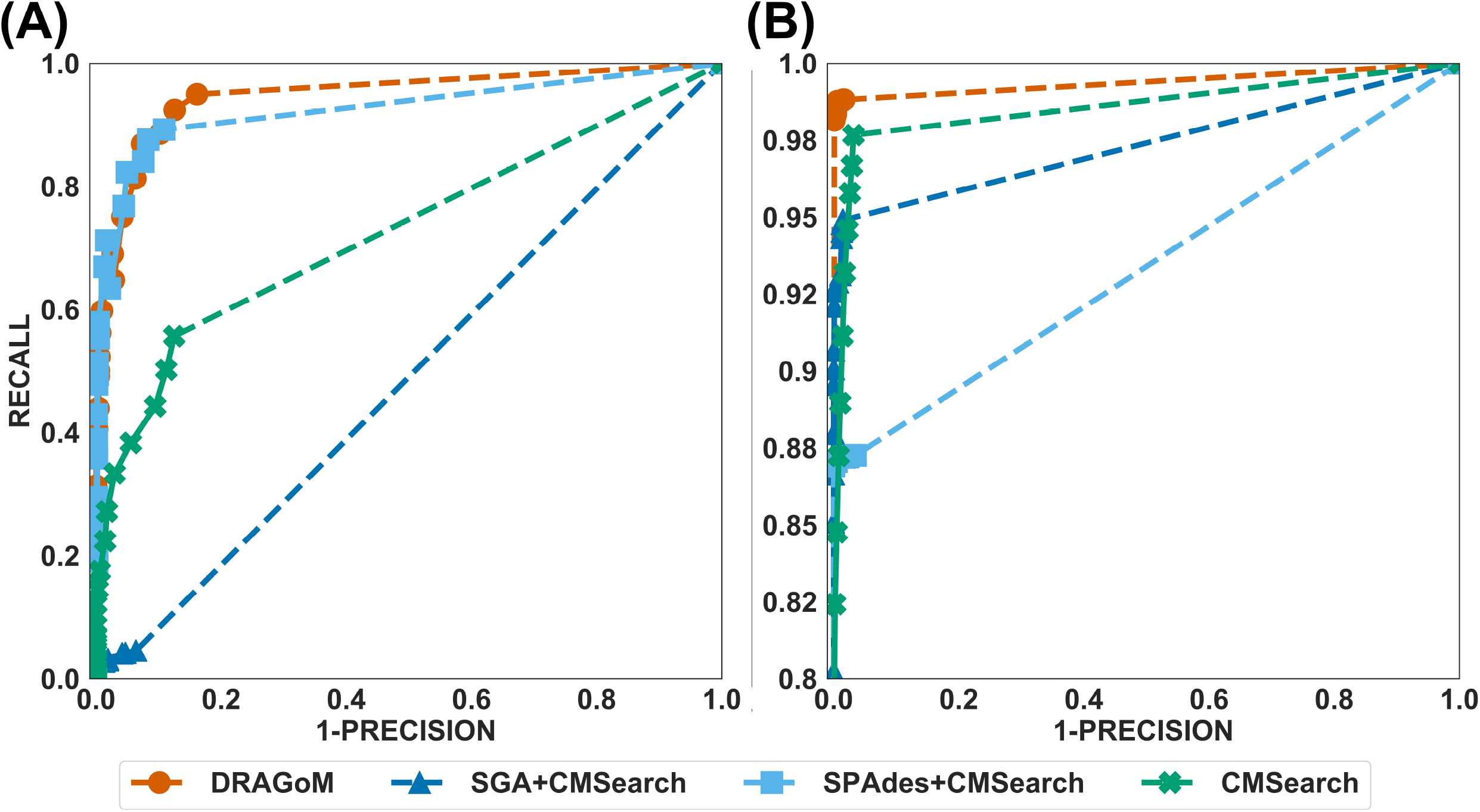
The ROC curves for searching (A) 93 non-16s rRNA ncRNA families and (B) 16s rRNA using the corresponding programs against DS3 (the simulated marine dataset).

DS5 (subsampled CAMI) was profiled using the largest number of querying ncRNA families (276); hence we categorize the performance of non-16s rRNA searches based on the ncRNA families’ sequence identity and average length (Figure 6 and Table 3). Although the performances differed in different categories of ncRNA families, DRAGoM consistently showed the best performance. The lowest performance gain made by DRAGoM was for the category with <50% sequence identity and 200-400bp length, showing a gain of 0.6% in F-score and 2.4% in AUC compared to the second-best method SPAdes+CMSearch (Figure 6(B)). The largest gain made by DRAGoM was found in the category with 50-70% sequence identity and 200-400bp length Interestingly, where the improvement was 11.4% in F-score (as compared to SPAdes+CMSearch) and 10.1% in AUC (as compared to CMSearch). Our interpretations of the difference in performance gain in different categories of ncRNA families are present in the Discussion section. For 16s rRNA, DRAGoM had the best performance F-score (96.4%, Table 4) but the second-best performance in AUC (96.8%, compared to the best performance of 97.6% made by CMSearch).

**Table 3.** Performance summary of the tested methods on DS5 (for non-16s rRNA queries). The highest performance of each category is bolded.

| Dataset | Matrices | DRAGoM | SGA+CMSearch | SPAdes+CMSearch | CMSearch |
| --- | --- | --- | --- | --- | --- |
| Identity: <50%<br>length: 100 - 200 | Precision | 77.5% | **89.2%** | 77.3% | 80.7% |
|  | Recall | **81.2%** | 32.6% | 74.3% | 50.3% |
|  | F1 | **79.3%** | 47.8% | 75.8% | 61.9% |
|  | AUC | **82.4%** | 61.7% | 78.9% | 66.4% |
| Identity: <50%<br>length: 200 - 400 | Precision | 71.3% | 73.0% | 72.1% | **81.4%** |
|  | Recall | **90.7%** | 35.2% | 87.9% | 39.5% |
|  | F1 | **79.8%** | 47.5% | 79.2% | 53.2% |
|  | AUC | **87.6%** | 52.0% | 85.2% | 62.2% |
| Identity: 50 - 70%<br>length: 100 - 200 | Precision | 85.4% | **86.6%** | 83.4% | 79.2% |
|  | Recall | **81.8%** | 40.4% | 76.1% | 78.1% |
|  | F1 | **83.6%** | 55.1% | 79.6% | 78.6% |
|  | AUC | **87.8%** | 65.4% | 82.3% | 80.7% |
| Identity: 50- 70%<br>length: 200 - 400 | Precision | 81.5% | **94.1%** | 78.0% | 87.3% |
|  | Recall | **87.2%** | 31.8% | 68.4% | 61.1% |
|  | F1 | **84.3%** | 47.5% | 72.9% | 71.9% |
|  | AUC | **85.3%** | 63.4% | 75.0% | 75.2% |
| Identity: 70 - 90%<br>length: 100 - 200 | Precision | 82.8% | **88.8%** | 85.7% | 77.4% |
|  | Recall | **85.7%** | 39.3% | 75.7% | 82.3% |
|  | F1 | **84.2%** | 54.5% | 80.4% | 79.8% |
|  | AUC | **87.5%** | 65.5% | 81.6% | 81.6% |
| Identity: 70 - 90%<br>length: 200 - 400 | Precision | 83.0% | **96.3%** | 88.5% | 83.6% |
|  | Recall | **89.1%** | 23.0% | 65.5% | 73.8% |
|  | F1 | **86.0%** | 37.1% | 75.3% | 78.4% |
|  | AUC | **87.5%** | 59.6% | 77.3% | 81.7% |
| Identity: > 90%<br>length: 100 - 200 | Precision | 74.2% | **94.6%** | 81.8% | 71.7% |
|  | Recall | **92.8%** | 23.6% | 81.5% | 85.9% |
|  | F1 | **82.5%** | 37.8% | 81.7% | 78.2% |
|  | AUC | **87.7%** | 59.4% | 83.5% | 82.3% |
| Identity: >90%<br>length: 200 - 400 | Precision | 90.1% | **99.7%** | 94.2% | 83.8% |
|  | Recall | **95.7%** | 17.7% | 84.9% | 72.3% |
|  | F1 | **92.8%** | 30.0% | 89.3% | 77.6% |
|  | AUC | **93.9%** | 58.7% | 89.5% | 81.6% |

**Table 4.** Performance summary of the tested methods on DS1-DS5 (for 16s rRNA queries). The highest performance of each category is bolded.

| Dataset | Matrices | DRAGoM | SGA+CMSearch | SPAdes+CMSearch | CMSearch |
| --- | --- | --- | --- | --- | --- |
| DS1 | Precision | 99.9% | 99.0% | **100.0%** | 97.1% |
|  | Recall | **99.2%** | 96.0% | 92.4% | 98.1% |
|  | F1 | **99.5%** | 97.5% | 96.0% | 97.6% |
|  | AUC | **99.6%** | 98.4% | 96.2% | 98.8% |
| DS2 | Precision | 96.2% | **98.7%** | 97.5% | 96.5% |
|  | Recall | **99.9%** | 99.7% | 97.4% | 99.5% |
|  | F1 | 98.1% | **99.2%** | 97.5% | 98.0% |
|  | AUC | 98.1% | **99.8%** | 97.5% | 99.4% |
| DS3 | Precision | 99.7% | 98.6% | **100.0%** | 96.8% |
|  | Recall | **98.6%** | 94.9% | 87.0% | 97.7% |
|  | F1 | **99.1%** | 96.7% | 93.0% | 97.2% |
|  | AUC | **99.3%** | 97.4% | 93.4% | 98.6% |
| DS4 | Precision | 91.9% | 97.2% | **97.7%** | 97.0% |
|  | Recall | **96.5%** | 95.0% | 89.8% | 79.8% |
|  | F1 | 94.2% | **96.1%** | 93.6% | 87.6% |
|  | AUC | 94.4% | **96.4%** | 94.1% | 88.3% |
| DS5 | Precision | **95.1%** | 94.7% | 94.6% | 94.2% |
|  | Recall | **97.7%** | 96.9% | 92.2% | 97.6% |
|  | F1 | **96.4%** | 95.8% | 93.4% | 95.9% |
|  | AUC | 96.8% | 96.6% | 94.1% | **97.6%** |

Taken together, DRAGoM consistently delivered superior search performance in nearly all datasets and all categories of querying ncRNA families. Specifically, DRAGoM produced the best ncRNA homology prediction for all non-16s rRNA in all datasets, and two out five datasets (DS1 and DS3) for 16s rRNA searches (DRAGoM was the second-best method for the other three cases). The assembly-based approach SPAdes+CMSearch seemed to be the second-best choice overall. However, the read-based approach CMSearch appeared to be the second-best choice when analyzing ncRNA families with sequence identity between 70-90% and length between 200-400bp (Figure 6(F)) and in the searches of 16s rRNAs on DS1, DS3, and DS5. Comparably, DRAGoM was the most robust method in addition to its superior performance.

### Analyzing a real case-control Type-2 diabetes dataset with DRAGoM

To showcase real-world applications of DRAGoM, and demonstrate the value of microbial ncRNA in characterizing human diseases, we used DRAGoM to profile 1,112 ncRNA families for all twelve samples in DS6 (metadata available from Supplementary Table 3). The datasets were first subject to standard quality trimming, followed by the DRAGoM homology search, and finally differential abundance significance calculation using DESeq2 [62] (details available in Supplementary Methods). Out of the total 1,112 ncRNA families searched, 342 of them had recruited at least one read in either of the twelve samples (see Supplementary Table 4). The top 50 ncRNA families with the most significant abundance difference were used as the potential biomarkers for the characterization of T2D from healthy controls.

Analysis of the top ncRNA families revealed the gut microbiome’s metabolic and physiological changes under the T2D condition. In general, we found that gene expression was more active in the microbes under the T2D condition, potentially due to their increased access to glucose and fatty acid in the environment as food sources. Specifically, we observed a reduced abundance of RF00032 (histone mRNA 3’UTR stemloop, *p*-value 3.6 × 10^−3^) in T2D samples, which is key to maintaining the chromatin structure. A decreased histone abundance level may suggest less packed chromosomes, which is often tied to more active gene transcription. We also observed a reduced abundance of RF00106 (RNAi, *p*-value 1 × 10^−2^) in T2D samples, which initiates the degradation of its target mRNA upon binding. A lower degradation activity would also imply a higher gene expression level. Moreover, the retrotransposition activity was also more active, as demonstrated by the higher abundances of group II intron families in under T2D (RF01998 with a *p*-value 1.3 × 10^−2^; RF02004 with a *p*-value 1.8 × 10^−2^; and RF02001 with a *p*-value 4.4 × 10^−2^). These ncRNA families are often found in transposons, plasmids, insertion sequence islands, and pathogenicity islands [63]. However, we do note that the datasets used here were metagenomic rather than metatranscriptomic. Hence, the abundance profile may not reflect the actual expression levels of those ncRNA families.

To further investigate whether these ncRNA families can be used as biomarkers to characterize T2D, we performed unsupervised hierarchical clustering (see Supplementary Methods) of the abundance profile (as RPKM) of the top 50 ncRNA families (Figure 7). All control samples were cluster together, indicating that they share the most similar ncRNA abundance profiles. Another four T2D samples (SRR341587, SRR341583, SRR341585, and SRR341624) were subsequently clustered with the healthy samples. The other two T2D samples (SRR341586 and SRR341588) seemed to have distinctive ncRNA abundance profiles relative to the other ten samples, as demonstrated by the ultra-high abundance of the ncRNA families shown in the upper half of the heatmap. Principal component analysis (PCA) was also performed on the top 50 ncRNAs’ abundances to verify the clustering results (Figure 8). Similar to the clustering result, the control samples were also well clustered, but the T2D samples exhibited a higher variance. To investigate the potential reasons why these three T2D samples had distinct ncRNA abundance profiles, we further analyzed the available metadata associated with the samples (Supplementary Table 3). We noted that SRR341586 had the smallest BMI (Body Mass Index) among all 12 samples (Z-score −2.42). SRR341587, on the other hand, was the tallest (Z-score 2.51) and the heaviest (Z-score 2.16) subject. SRR341588 had the lowest total cholesterol level (Z-score −2.37). Whether the distinct ncRNA abundance profile is due to the above sample selection bias remains unclear and was subject to experimental investigation. Another explanation could be that these T2D patients corresponded to different T2D subtypes [64] or were in different developmental stages of T2D.

## DISCUSSION

We have demonstrated using benchmark data that DRAGoM can improve ncRNA homology search in comparison to traditional read-based and assembly-based strategies. In addition to the higher performance, another unique advantage of DRAGoM is its robustness. We observed from the benchmark results that the homology search performance is both query-dependent and dataset-dependent. For example, in DS5 (CAMI), SPAdes+CMSearch performed better than CMSearch when searching ncRNA families with identity <50% and between 100-200bp long (Figure 6(A)), but worse when searching ncRNA families with identity 70-90% with the same length range (Figure 6(E)). We conjecture that some factors could have contributed to such differences. If the ncRNA families are highly divergent, sequence information alone may not be sufficient for its detection. Therefore, the complete secondary structure information needs to be reconstructed for its detection. It may explain the higher performance of SPAdes+CMSearch when searching low-identify families (Figure 6(A)). On the other hand, reads from highly conserved ncRNA families are more easily detected, but may be treated as repeats and be deleted/collapsed during assembly. This may explain the higher performance of CMSearch when searching high-identity families (Figure 6(E)). Moreover, the homology search performance was also dataset-dependent. For example, CMSearch performed better than SGA+CMSearch in the 16s rRNA search against DS3 (Figure 4(B)), but worse against DS4 (Figure 5(B)). The assembly-based homology search was more recalcitrant to experimental noise (e.g., the experimentally generated DS4), as the homology information loss of a given read due to sequencing error may be rescued by its adjacent overlapping reads. This may explain the higher performance of SGA+CMSearch when searching 16s rRNA in DS4. Nevertheless, no method demonstrated a consistently-high performance in all queries and datasets, except DRAGoM. This robustness is another advantage of DRAGoM in addition to its higher performance.

**Figure 5:**
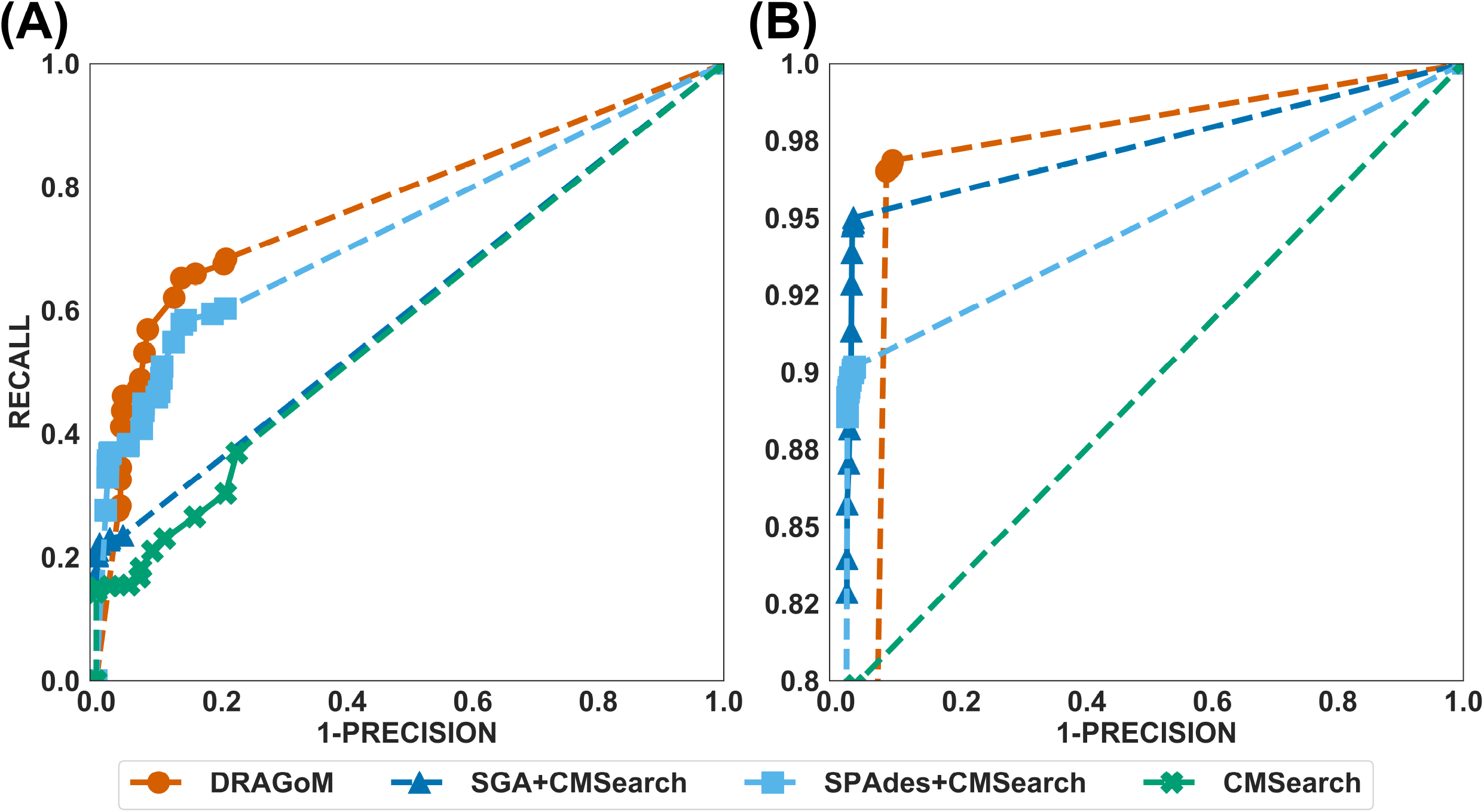
The ROC curves for searching (A) 60 non-16s rRNA ncRNA families and (B) 16s rRNA using the corresponding programs against DS4 (the subsampled human gut dataset).

**Figure 6:**
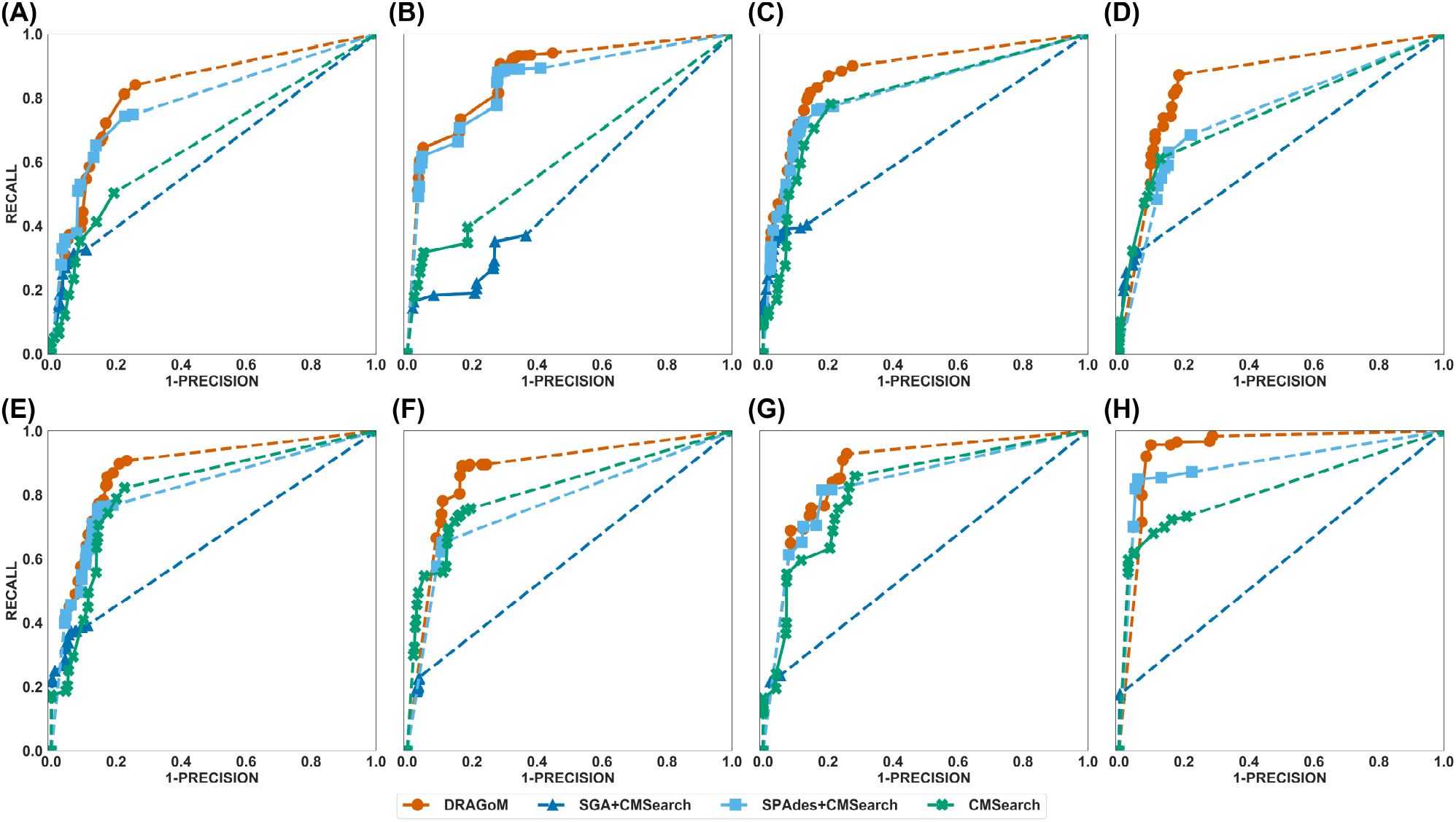
The ROC curves for searching (A) 276 non-16s rRNA ncRNA families and (B) 16s rRNA using the corresponding programs against DS5 (the subsampled CAMI dataset).

**Figure 7:**
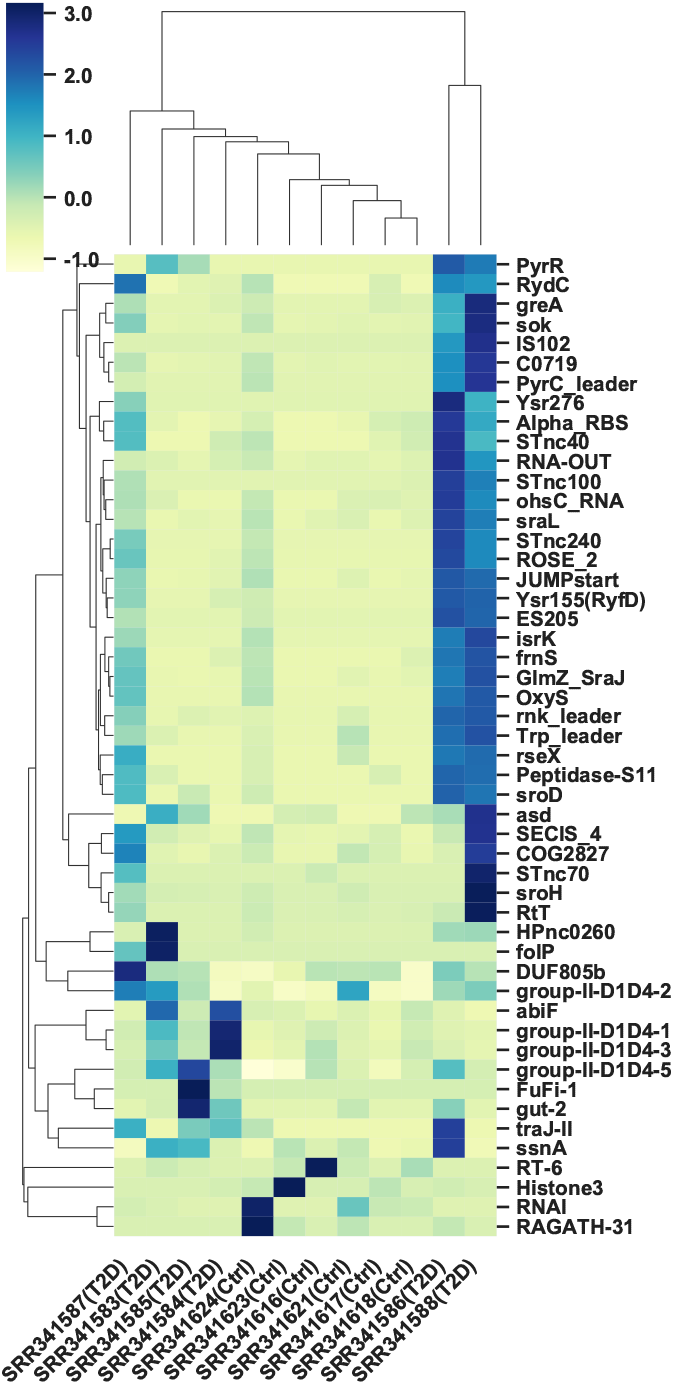
The clustering heatmap for the top 50 differentially abundant ncRNA families in DS6 (6 healthy controls and 6 T2D cases).

**Figure 8:**
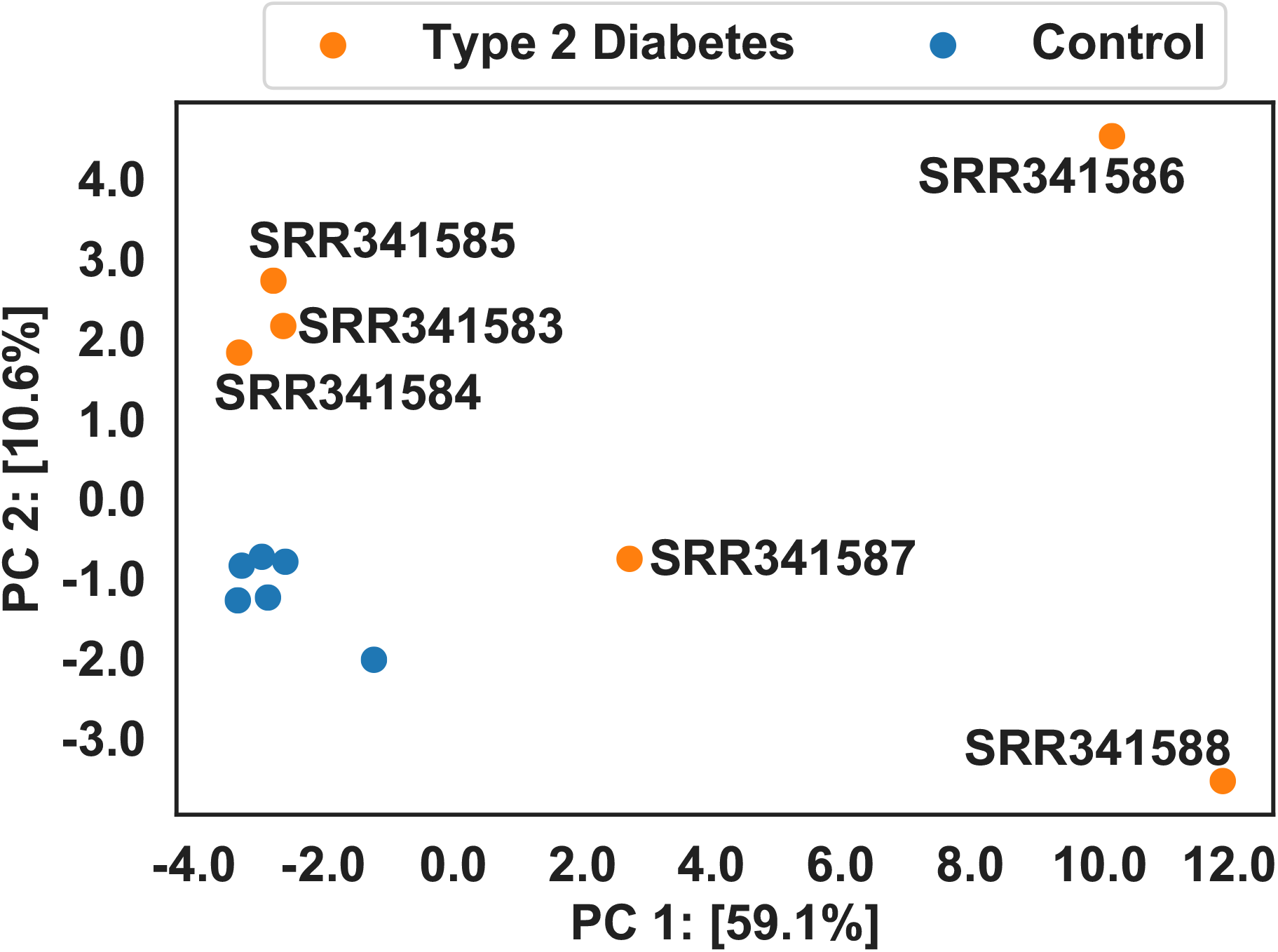
The PCA plot for the top 50 differentially abundant ncRNA families in DS6 (6 healthy controls and 6 T2D cases).

Because DRAGoM directly operates on an assembly graph, the quality of the assembly graph will likely affect the performance of DRAGoM. Currently, string graph and de Bruijn graph dominate the modeling of sequence overlap information in de novo assembly. DRAGoM, which is based on the combination of the two graphical models, outperformed the use of either of them alone (i.e., SGA+CMSearch and SPAdes+CMSearch). The observation is consistent with our current understanding of the two models, where each of them has its unique advantages (where the string graph accurately represents the intact information and the de Bruijn graph generates more contiguous and complete assembly).

SPAdes+CMSearch outperformed SGA+CMSearch in most cases we have observed, suggesting that the reconstruction of complete secondary structure is more important than the preservation of polymorphism information in ncRNA homology search. However, our conjecture is by no means conclusive, as shown by cases where SGA+CMSearch outperformed SPAdes+CMSearch (e.g., Figure 5(B)).

We expect to further improve the speed of DRAGoM in the future. Specifically, the efficiency bottleneck of DRAGoM comes from the fact that it needs to exhaustively align the querying CM with all paths generated from anchors. We envision two potential ways to improve DRAGoM’s speed, though improved path filtering heuristics and graph simplification techniques. We plan to filter out paths that have inconsistent GC content, coverage, and incompatible covariant mutations before the CM alignment. We also expect to reduce the complexity of the assembly graph through incorporating additional information, such as paired-end, long-read, or Hi-C data if applicable [65]. DRAGoM was slower when searching long ncRNA families, as the time for CM alignment and the number of candidate paths to align both grow with the length. As a result, for a long querying ncRNA family, we plan to break it down into a set of smaller components by temporarily removing long-range interactions, align each small component individually, and check if the removed long-range interactions can be recovered given the alignments. This heuristic has been proven effective in speeding up the alignment of RNA structural motifs with pseudo-knots while retaining satisfying alignment quality [66]. We anticipate significantly reduced running time of DRAGoM with the above heuristics.

In addition to the ncRNA family abundance profile, DRAGoM may also be used to improve the taxonomic analysis of metagenomic datasets in two ways. First, DRAGoM can improve the traditional 16s rRNA-based taxonomic analysis. The existing methods first identify 16s rRNA reads from the metagenomic datasets, perform local assembly on the 16s rRNA reads, and then infer the taxonomy [24]. DRAGoM can improve this strategy in its first step via providing a more complete and reliable set of 16s rRNA reads. The second way that DRAGoM may improve taxonomic analysis is through facilitating the use of ncRNA families as taxonomic biomarkers, in a similar way as the protein taxonomic biomarkers [34, 67, 68]. However, the current implementation of DRAGoM does not output complete ncRNA genes, because many homologous paths appeared to be redundant. We plan to incorporate a more sophisticated algorithm into DRAGoM to infer the non-redundant set of homolog paths. One possible way would be via finding the minimum set of paths that cover the entire homolog-read assembly graph (as in REAGO [24]). Another way would be using the similar statistical methods for isoform prediction and quantification in RNA-seq data analysis [69]. We believe that the detection of novel ncRNA taxonomic biomarkers may help us to obtain unbiased and comprehensive taxonomic profiles.

Although we have shown the potential of DRAGoM in identifying ncRNA biomarkers for characterizing T2D gut microbiome, the study bears potential limitations. First, the sample size (six cases vs. six controls) was not large enough to draw reliable conclusions regarding the ncRNA biomarkers’ predictive power, and we expect to include more datasets in the future. Second, all selected samples were from the female Chinese Han population, and the discoveries had not been validated on other populations. Therefore, the identified ncRNA biomarkers could resulted from overfitting and may not extend to other populations. Third, the functional analysis and interpretation of the biomarkers were purely based on the existing functional annotation, and were subject to experimental validation. Some of the most differentially-abundant ncRNA families (e.g., RF00124, which ranked the top in our analysis) had no functional annotation either; therefore, they were not included in our functional analysis. We believe this issue will be alleviated as more experimental evidence is being accumulated and incorporated into ncRNA functional databases. Finally, the ncRNA family abundance may not correspond to their final expression levels, as the selected datasets were metagenomic instead of metatranscriptomic. We note that metatranscriptomic data could provide more direct insights into the physiological status of the microbiome, and could be a better technology for understanding ncRNAs function [70]. We are current working on extending DRAGoM to metatranscriptomic datasets.

Finally, once the speed up of DRAGoM is fulfilled, we plan to apply it to identify more ncRNA biomarkers for more disease types and couple them with the existing protein biomarker for more accurate diagnoses. We plan to disseminate the identified biomarkers for different diseases via publicly available databases, which will have a great value for designing cost-effective early-diagnostic approaches based on amplicon sequencing or microarrays. In summary, we have developed DRAGoM, a novel algorithm for family-based ncRNA homology search against metagenomic sequencing data. We have demonstrated the advantages of DRAGoM as compared to the traditional read-based and assembly-based approaches, and its potential in bacterial ncRNA biomarker discovery for disease characterization using a real case-control study of T2D. DRAGoM is implemented using GNU C++ and Python, and is freely available from https://github.com/benliu5085/DRAGoM under the Creative Commons BY-NC-ND 4.0 License Agreement (https://creativecommons.org/licenses/by-nc-nd/4.0/).

## Acknowledgement

The authors would like to thank Mr. Hao Xuan and Mr. Adam Podgorny for their contributions in software testing and manuscript draft proofreading. This work is funded by the National Science Foundation EPSCoR First Awards in Microbiome Research and the National Science Foundation CAREER award DBI-1943291.

## Notes

https://cbb.ittc.ku.edu/DRAGoM.html

https://github.com/benliu5085/DRAGoM

### Competing Interest Statement

The authors have declared no competing interest.

