## Supplementary Methods for "Metagenomic Noncoding RNA Profiling and Biomarker Discovery"

### Supplementary Methods for "Metagenomic Non-coding RNA Profiling and Biomarker Discovery with DRAGoM"

#### The DRAGoM Workflow

DRAGoM was developed in C++ (g++ v 4.8.5) and python (2.7), containing the following stages.

##### Stage 1: Generating SPAdes Contigs

We generated SPAdes [1] (v3.13.0) contigs using the following script:

```
spades.py --meta -t 16 -m 1024 \
-1 read1.fq -2 read2.fq \
-o spades
```

For DS3, where the input reads were interleaved paired-end reads, the command line we ran was:

```
spades.py --meta -t 16 -m 1024 \
--12 read.fq \
-o spades
```

##### Stage 2: Generating SGA String Graph

We generated the SGA [2] (0.10.15) overlap graph using the following steps (as recommended by SGA best practice):

(a) preprocess

```
sga preprocess -o reads.pp.fastq --pe-mode 1 read1.fq read2.fq
sga preprocess -o reads.pp.fastq --pe-mode 2 read.fq
```

(for DS3, whose paired-end reads are stored in a single file)

(b) error-correction

```
sga index --no-reverse -t 16 reads.pp.fastq
sga correct -k 55 --learn -d 256 -t 16 reads.pp.fasta
```

(c) remove duplicate reads

```
sga index -t 16 reads.pp.ec.fa
sga rmdup -t 16 reads.pp.ec.fa
```

(d) building overlap graph

```
sga overlap -t 16 -m 30 reads.pp.ec.rmdup.fa
```

Once the overlap graph was generated, we developed an in-house script to convert the overlap graph into a string graph. The source code of this conversion script is available from the GitHub repository of DRAGoM: https://github.com/benliu5085/DRAGoM.

##### Stage 3: Generating the Hybrid Graph

Define *terminal* as an edge (i.e., unitig) in the string graph that has either of its ends has a degree of 0; also define *open end* as the terminal end that has 0 degree. For SPAdes contigs, we further validated them by aligning the individual reads back and only retained the ones that have no coverage hole (i.e., each position in the contig is covered by at least one read). We called these contigs as the *trusted contigs*. We then align the terminals against the trusted SPAdes contigs using BWA (0.7.16a-r1185-dirty) [3]:

```
bwa index spades.contigs.fasta
bwa mem -t 16 -a -T 45 spades.contigs.fasta og.fq > og.sam
```

By using -T 45, we only considered alignments with scores >=45 (per BWA manual, +1 was issued for a match, -4 for a mismatch, and -6 for a gap). We further filter the alignments by ensuring no clipping on the open-end side of the terminal sequence and retaining only alignments that are longer than 100bp. Note that the terminal edges correspond to condensed paths in the string graph, rather than an overlap between two reads as in a traditional overlap graph. In this case, our cutoff of 100bp still applies to datasets with an average read length less than 100bp. Once the alignments were collected, we then connect two terminals if they can be aligned to the same SPAdes contig, using the sequence indicated by the corresponding SPAdes contig interval. If the terminal is aligned to a SPAdes contig with no other alignment, the terminal is extended based on the corresponding prefix/suffix sequence of the SPAdes contig. Finally, the SPAdes contigs recruited no alignment were further retained as isolated vertices in the hybrid graph.

**Stage 4: Anchoring**

For a given ncRNA query, we used CMsearch (INFERNAL [4] 1.1.2) to search the query against each edge of the resulted hybrid graph:

```
cmsearch -T t1 --nohmmonly --rfam --cpu 16 \
    --tblout out.tblout RNA.cm hybrid.graph
```

All hits reported by CMSearch under its default cutoff were used as anchors. “-T t1” represents the use of family-dependent cutoffs as included in the CMs.

**Stage 5: Generating candidate paths**

For a given ncRNA query, we prioritized the anchors based on their CMSearch E-values. The anchors were then extended towards both directions using a breath-first-search (BFS) algorithm. The extension length is at most 120% of the total length of the prefix or suffix of the querying ncRNA family (based on the anchor position in the ncRNA family), or 100, whichever is smaller. During the BFS, the traversed edges were masked out and no longer available for the lower-priority anchors. Finally, the resulted paths were clustered using CD-HIT[5] (4.7) to remove redundancy:

```
cd-hit-est -M 0 -c 0.99 -T 16 -o path.cd.fa -i path.fa
```

**Stage 6: Predicting homologous ncRNA reads**

To identify homologous ncRNA paths, we searched the ncRNA query against the candidate paths using CMSearch (INFERNAL [4] 1.1.2)

```
cmsearch --cut_ga --nohmmonly --rfam --cpu 16 \
    --tblout out.tblout RNA.cm path.cd.fa
```

We collected the output of the above search as the set of homologous paths (with unaligned flanking sequences trimmed).

As the homologous path set could be large, we performed a two-stage CD-HIT redundancy removal step. In the first stage, redundant (>99% sequence identity) homologous paths are removed. In the second stage, the remaining homolog paths are clustered with 90% sequence identity to avoid missing read mappings due to excessive multi-mapping. Specifically, we ran

```
cd-hit-est -M 0 -c 0.9 -T 16 -o 90.RNA.cd.fa -i RNA.cd.fa
```

We then partition the entire homologous path set based on the resulted clustering, such that within each partition no more than 3,000 sequences are from the same cluster. Finally, we aligned the original reads against these partitions using BWA (0.7.16a-r1185-dirty) [3]:

```
bwa index RNA.fa
bwa mem -t 16 -a RNA.fa read.fa
```

We considered reads that can be aligned to any homologous path with >60% of its total length as the final set of homologous ncRNA reads.

#### Experimental Materials

**DS1 (REAGO)**
The microbial genomes of this simulated dataset were adopted from REAGO [6]. The reference genomes and their relative abundances and accumulated coverages can be found from Supplementary Table 1. The reads were simulated using WGSIM [7]:

```
wgsim -N2326963 -1100 -2100 -d180 -S7 -e0.01 -r0 \
      stagger.reago.fna reago.read1.fq reago.read2.fq
```

**DS2 (simulated streptococcal genomes)**
This dataset was generated using 8 *Streptococcus* genomes to investigate cases where highly-related genomes are present. The reference genomes and their relative abundances and accumulated coverages can be found from Supplementary Table 1. The reads were simulated using WGSIM [7]:

```
wgsim -N300000 -1100 -2100 -d180 -S7 -e0.01 -r0 \
      strep.fna strep.read1.fq strep.read2.fq
```

**DS3 (simulated marine)**
This simulated data composed of typical microbiomes found in marine. The reference genomes and their relative abundances and accumulated coverages can be found from Supplementary Table 1. The reads were generated using WGSIM [7]:

```
wgsim -N1850000 -1100 -2100 -d180 -S7 -e0.01 -r0 \
      stagger.reago.fna reago.read1.fq reago.read2.fq
```

**DS4 (subsampled human gut microbiome)**
This dataset corresponded to a real human gut metagenomic sequencing dataset, and was downloaded from SRA (SRR341583). The raw reads were first quality-trimmed using Trimmomatic [8] (v0.38):

```
java -jar trimmomatic-0.38.jar \
      PE read1.fastq read2.fastq \
      forward.paired.fq.gz forward.unpaired.fq.gz \
      reverse.paired.fq.gz reverse.unpaired.fq.gz \
      ILLUMINACLIP:TruSeq2-PE.fa:2:30:10 \
      LEADING:3 \
      TRAILING:3 \
      SLIDINGWINDOW:4:15 \
      MINLEN:45 -threads 16;
```

After trimming, only the reads that remained paired after trimming were retained. The reads were then mapped (using BWA) to a set of reference genomes that were often found in the human gut environment (Supplementary Table 1).

```
bwa index SRR341583.fna
bwa mem -t 16 SRR341583.fna forward.paired.fq reverse.paired.fq
```

Paired-end reads that has at least one of the ends mapped to the reference genomes were retained, resulted in a total of 11,228,362 reads.

**DS5 (subsampled CAMI)**
This simulated data was down-sampled from the CAMI [9] toy test dataset labeled with "Medium_Complexity". The reference sequences (genome, scaffold, and contig) provided by CAMI with a designed abundance lower than 10 were selected for the subsampling process (Supplementary Table 1). The reads were quality trimmed using Trimmomatic [8] (v0.38), mapped against the reference genomes using BWA, and subsampled in the same way as DS4. The final dataset contained 31,311,294 read pairs.

**DS6 (the case-control T2D dataset)**
This dataset was obtained from a study of human T2D-associated gut microbiome [10]. Six datasets of healthy controls (SRR341616, SRR341617, SRR341618, SRR341621, SRR341623, SRR341624) and six datasets of T2D cases (SRR341583, SRR341584, SRR341585, SRR341586, SRR341587, SRR341588) were selected. To eliminate confounding factors, all samples were from females. All datasets were subject to quality trimming in the same way as DS4. The number of reads contained in each dataset is summarized in Supplementary Table 1.

**Dataset Availability**
All datasets generated in-house (i.e., DS1-5) can be downloaded from https://cbb.ittc.ku.edu/DRAGoM.html. DS6 can be downloaded from the SRA database with the accession numbers provided above.
